# Normalizing LZ+ MYPT1 Expression Prevents the Development of HFpEF

**DOI:** 10.64898/2026.08.19.745871

**Authors:** Young Soo Han, Teresa M Pfiefer, Bin Zhang, Matthew J Fogarty, Gary C. Sieck, Frank V Brozovich

## Abstract

**Background:** Heart failure (HF) is classified by ejection fraction: reduced EF (<40%) is HFrEF and preserved EF (>50%) is HFpEF. Unlike HFrEF, no therapeutic agent improves mortality in HFpEF. The molecular mechanism that produces HFpEF is not completely understood, but the cascade of pathology that produces HFpEF is thought to begin with changes in vascular reactivity, including a decrease in NO mediated vasodilatation, which coupled with subsequent changes in contractility, energetics and coronary blood flow produce HFpEF. If abnormal vascular reactivity is the initial step in the pathological cascade that produces HFpEF, restoring and/or improving vascular reactivity could represent a novel treatment strategy. Vascular reactivity is primarily regulated by myosin light chain phosphatase, which has catalytic, myosin targeting (MYPT1) and 20kDa subunits. Alternative mRNA splicing of exon24 (E24) of the MYPT1 transcript produces MYPT1 isoforms that differ by the presence or absence of a COOH-terminal leucine zipper (LZ+/LZ-); E24 exclusion produces an NO responsive LZ+ MYPT1, while E24 inclusion produces an NO unresponsive LZ- MYPT.

**Methods:** We used the mouse two-hit model of HFpEF (high fat diet and L-NAME) and treated mice with an anti-sense octo-guanidine targeting the 5’ splice site of E24 (ASO-E24) to increase the expression of the NO responsive, LZ+ MYPT1 isoform in vascular smooth muscle. Invasive and noninvasive hemodynamics were used to determine LV function.

**Results:** Compared to mice with HFpEF, ASO-E24 treatment maintains LZ+ MYPT1 expression (4.7±0.7au v 1.0±0.4au v 2.0±0.4au, control v HFpEF v ASO-E24 Rx, p<0.05), improves diastolic function; LVEDP (10±1mmHg v 20±4mmHg v 14±3mmHg, p<0.05), dP/dtmin (−8000±300mmHg/s v 6000±500mmHg/s v 8500±700mmHg/s, p<0.05), both early (E; 0.60±0.05m/s v 0.42±0.06m/s v 0.64±0.06m/s, p<0.05) and late diastolic filling (A; 0.38±0.03m/s v 0.24±0.02m/s v 0.47±0.04m/s, p<0.050 and also prevents the increase in lung weight (167±5g v 175±7g v 166±5g, p<0.05). Further, mice treated with ASO-E24 maintained normal relaxation to 8Br-cGMP (65±5% v 44±9% v 72±9%, p=0.05).

**Conclusion:** These data demonstrate that maintaining normal LZ+ MYPT1 expression and vascular reactivity prevent the development of HFpEF. These results are consistent with the hypothesis that abnormal vascular reactivity is the initial and primary step in the pathological cascade that produces HFpEF and ASO-E24, which is designed to preserve normal LZ+ MYPT1 expression and vascular reactivity, could represent a novel and effective treatment strategy for HFpEF.

**Clinical Prospective:** *What is New?:*

- In HFpEF, an anti-sense octo-guanidine targeting the 5’ splice site of exon 24 (ASO-E24) of the MYPT1 transcript increases the expression of the NO responsive, LZ+ MYPT1 isoform in vascular smooth muscle.
- Preserving LZ+ MYPT1 expression and normal vascular reactivity prevents the development of HFpEF.
- A decrease in LZ+ MYPT1 expression contributes to the development of HFpEF.

*What are the Clinical Implications?:*

- The results of the study demonstrate that abnormal vascular reactivity is an important step in the pathological cascade that produces HFpEF
- Increasing LZ+ MYPT1 expression could represent a novel treatment strategy for HFpEF.

## Introduction

Heart failure (HF) is classified by ejection fraction (EF); EF<40% is HFrEF and EF>50% is HFpEF. For HFrEF, guideline-directed medical therapy improves both symptoms and mortality ^1^. However, for HFpEF, no treatment has been demonstrated to reduce mortality ^2–9^. HFpEF is associated with diastolic dysfunction ^10–12^ and abnormal vascular reactivity including a resting vasoconstriction and reduced sensitivity of the vasculature to NO mediated vasodilatation ^12^. The molecular mechanism underlying HFpEF is unknown and recent work suggests that HFpEF is the result of a cascade of pathological processes beginning with reduced sensitivity to NO mediated vasodilatation and increased vascular tone/stiffness, which then combined with subsequent changes in cardiac contractility and energetics ultimately contribute to the development of HFpEF ^13,14^.

Vascular tone is regulated by the phosphorylation of the regulatory light chain (RLC) of smooth muscle (SM) myosin, which is controlled by the activity of myosin light chain (MLC) kinase and MLC phosphatase ^15^. MLC kinase is regulated by Ca^2+^-calmodulin ^16,17^, but the vast majority of signaling pathways that control vascular tone converge on MLC phosphatase ^15,18,19^. MLC phosphatase is a trimeric enzyme consisting of catalytic, myosin targeting (MYPT1) and 20kDa subunits ^15,18^. Alternative mRNA splicing of exon24 (E24) of the MYPT1 transcript produces MYPT1 isoforms that differ by the presence or absence of a COOH-terminal leucine zipper (LZ+/LZ-); E24 exclusion (E24-) produces an NO responsive LZ+ MYPT1, whereas E24 inclusion (E24+) produces an NO unresponsive LZ- MYPT1 ^20,21^; LZ+/LZ- MYPT1 isoform expression regulates the sensitivity to NO mediated vasodilation ^22–25^. Thus, a decrease in LZ+ MYPT1 expression would reduce the sensitivity of the vasculature to NO ^20,22–26^ and lead to abnormal vascular reactivity ^24,25,27–29^.

We have previously demonstrated that LZ+ MYPT1 expression is lower in rodents ^30^ and humans ^31^ with HFpEF, consistent with the hypothesis that abnormal NO mediated vasodilatation is the first step in the pathological cascade that produces HFpEF ^13,14^. If this is the case, interrupting the pathological cascade at its initial step, or a decrease in LZ+ MYPT1 expression, and maintaining normal vascular reactivity could prevent the subsequent changes in cardiac contractility and energetics, which potentially represent a novel treatment paradigm for HFpEF.

Recently, IP injection of an anti-sense octo-guanidine targeting the 5’ splice site of MYPT1 transcript has been demonstrated to suppress E24 inclusion of the MYPT1 transcript and increase LZ+ MYPT1 expression from day 1 to at least 28 days after injection ^32^. The present study was designed to test the hypothesis that maintaining LZ+ MYPT1 expression would prevent the development of HFpEF. To test this hypothesis, we used the mouse two-hit model of HFpEF ^33–35^ and examined whether suppressing E24 inclusion in the MYPT1 transcript and thereby maintaining normal LZ+ MYPT1 expression would prevent the development of HFpEF.

## Methods

### Animals

The Institutional Animal Care and Use Committee of the Mayo Clinic approved all experimental protocols, which conformed to the guidelines of the National Institutes of Health.

To suppress E24 inclusion, we administered an intraperitoneal (IP) injection of an anti-sense octo-guanidine targeting the 5’ splice site of MYPT1 transcript (ASO-E24, GeneTools, LLC, ASOMYPT130MER, Philomath, OR). The sequence of ASO-E24 and its target sequence at the 5’ splice site is shown in Figure 1. IP injection of an ASO targeting the 5’ splice site of the MYPT1 transcript has been demonstrated to both suppress E24 exon inclusion and increase the expression of the LZ+ MYPT1 isoform ^32^. For our experiments, 12-week-old C57BL6 mice (male) were obtained from The Jackson Laboratory. HFpEF was induced using the two-hit model, as previously described ^30,33^ ; mice were fed a high fat diet (HFD; Tekla TD 06414) and L-N^6^-nitro arginine methyl ester (L-NAME, 0.5g/l) in the drinking water for 4 weeks, and these mice were designated as HFpEF. In another groups of mice, mice were fed the high fat diet and L-NAME in the drinking water and on day 1, concomitantly treated with ASO-E24 (3 doses qod; 12.5mg/kg, 12.5mg/kg, 9.25mg/kg), and these mice were designated ASO-E24 Rx. The final group of mice received standard chow and water and served as normal controls (Control). Animals were sacrificed after 4 weeks of treatment (16 weeks of age), and tissues were collected for subsequent analysis. We did not include female mice in this study, because they do not develop HFpEF in response to HFD and L-NAME ^34,35^.

**Figure 1:**
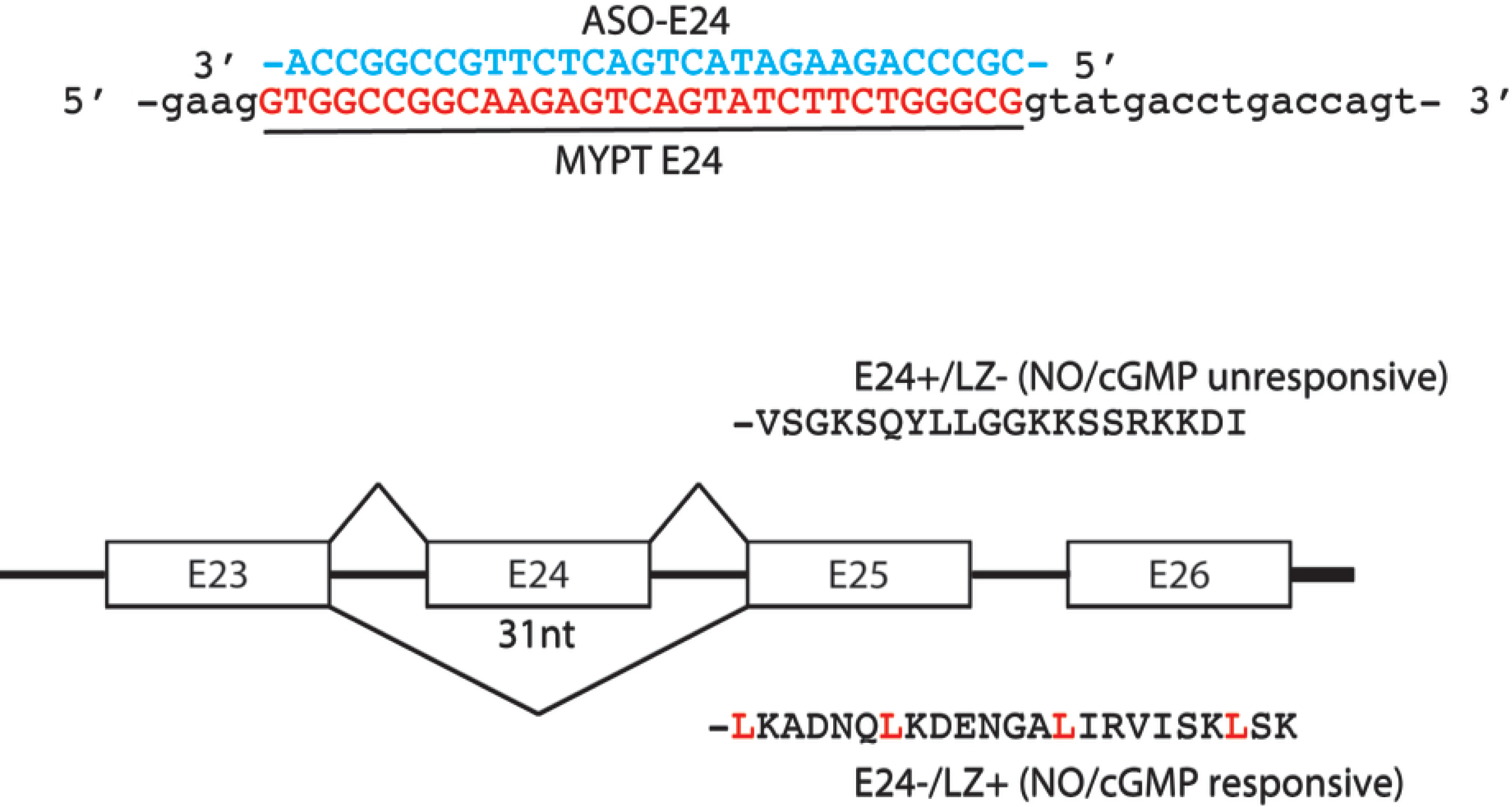
The sequence of ASO-E24 and its target sequence at the 5’ splice site. Sequence of ASO-E24 (blue) and its target sequence at 5’ MYPT1 splice site (red). Exclusion of E24 generates mRNA that codes for a COOH-terminal LZ motif (LZ+), while inclusion of E24 shifts the reading frame and codes for a premature stop codon and COOH-terminus that lacks the LZ motif (LZ-). Leucine residues are highlighted in red. The LZ+ MYPT1 isoform is required for PKG mediated activation of MLC phosphatase and is responsive to NO.

### RNA Analysis

Analysis of RNA was performed as previously described ^20,32,36^. Briefly, mesenteric vessels were dissected, cleaned and homogenized. Total RNA was purified with RNeasy® Mini Kit (QIAGEN Cat# 74104) and yield was quantified by optical absorbance (NanoDrop). RNA was reverse transcribed with Superscript reverse transcriptase enzyme and oligo dT primers using T100™ Thermal Cycle (Bio-Rad); 5’’-TGCAGTTGGAAAAGGCTACC-3’ (forward) and 5’-TCAAGGCTCCATTTTCATCC-3’ (reverse). MYPT1 exon24 splice variant PCR products were separated with 2.5% agarose gel electrophoresis and quantified using ImageLab software. E24 inclusion was quantified as the band densities representing E24 inclusion/(E24 inclusion + E24 exclusion).

### Immunoblotting

Immunoblotting was used to determine protein expression. As previously described ^29,30,37–41^, mesenteric vessels were homogenized in SDS sample buffer and total protein (TP) was resolved by SDS-PAGE, with sample loading normalized to TP within the band calculated from stain free precast gels (Bio-Rad Cat#64551870) ^30,37,38,42,43^. After SDS-PAGE, proteins were transferred to a Trans-Blot Turbo pack (Bio-Rad Cat# 1704157) in the Trans-Blot Turbo transfer system (Bio-Rad) and anti-MYPT1 (Abcam, ab32519) and anti-MYPT1 (Abcam, ab32519) and anti-LZ+MYPT1 ^41^ antibodies were used to visualize proteins. Left ventricular immunoblots were prepared in a similar manner and anti-SERCA2 (ab1137020, Abcam), anti-phospholamban (PLN, ab2199626, Abcam) and anti-S16 phospho-PLN (ab15000, Abcam), anti-troponin I (TnI, ab2199625, Abcam) and anti-S23/24 phospho-TnI (4004S, Cell Signaling) antibodies were used to visualize proteins. MyBP-C was visualized with SYPRO Ruby staining and phosphorylation of MyBP-C was defined using Pro-Q Diamond phosphoprotein staining, as described previously ^30,38^. The immunoblots were then scanned and analyzed using ImageLab software, and protein expression was normalized for total protein ^30,37,38,42,43^.

### 2.4. Invasive and Noninvasive Hemodynamics

As previously described, LV hemodynamics were obtained using a Millar catheter ^29,30^, and for noninvasive studies ^30^, transthoracic echocardiography was performed with the Vivid E9 ultrasound system (GE Healthcare).

### Vascular Reactivity

Mechanical studies were performed using previously published protocols ^37,44^. Mesenteric vessels were mounted on a DMT 4-channel myograph system ^45,46^ using 40µm stainless steel wire and stretched to Lo (length for maximal force). Preparations were allowed to equilibrate for 1h in continuously oxygenated physiological saline (PSS (mM); 140 NaCl, 3.7 KCl, 2.5 CaCl_2_, 0.8 MgSO_4_, 1.2 KH_2_PO_4_, 0,03 EDTA, 5.5 glucose, 25 HEPES, pH 7.4). Vessels were stimulated to contract with 80 mM KCl depolarization and after force reached a steady state, relaxation to the cell permeable cGMP analog, 8Br-cGMP, was measured.

### Histology

Cardiac hypertrophy was assessed from hematoxylin and eosin (H&E) stained LV tissue sections as previously described ^30,37^. Cardiomyocyte cross-sectional area was measured using ImageJ software (V1.48; National Institutes of Health, Bethesda, MD) as described^30,37^.

### Statistics

All data are presented as mean ± SEM (n refers to the number of animals). Prior to experimentation, power analysis was performed to determine the minimum number of animals required to detect biologically meaningful differences between groups. Using pilot data and prior publications to estimate endpoint-specific standard deviations (Z, varying by measurement), a Type I error rate of α = 0.05, and desired power of 0.8, we determined that n = 6 animals per group would provide sufficient power to detect a 20% difference in the primary outcome measures. Group comparisons were performed using an ANOVA. When significant differences were found, two-tailed Student’s t-tests were used for post hoc comparisons. Statistical significance was defined as p < 0.05. All analyses were conducted using GraphPad Prism (V11) or equivalent statistical software.

## Results

### Invasive Hemodynamics

Invasive hemodynamics (Fig 2) were used to assess LV function in control (n=4), HFpEF (n=4) and ASO-E24 treated (n=4) mice (Table 1). HFpEF mice are hypertensive compared to control mice (140±5/93±4mmHg v 94±3/59±2mmHg, p=0.0001). Blood pressure in the ASO treated mice is also elevated compared to normal control mice (130±10/75±7mmHg, p=0.0222), but not different compared to the mice with HFpEF (p>0.05). There was no difference in dP/dtmax, but dP/dtmin is lower in HFpEF (−6000±500mmHg/s) compared to both control (−8000±300mmHg/s, p=0.008) and ASO treated mice (−8500±700mmHg/s, p=0.0142). Further LVEDP was elevated in HFpEF compared to control mice (20±4mmHg v 10±1mmHg, p=0.022), and LVEDP was similar in control and ASO-E24 treated mice.

**Figure 2:**
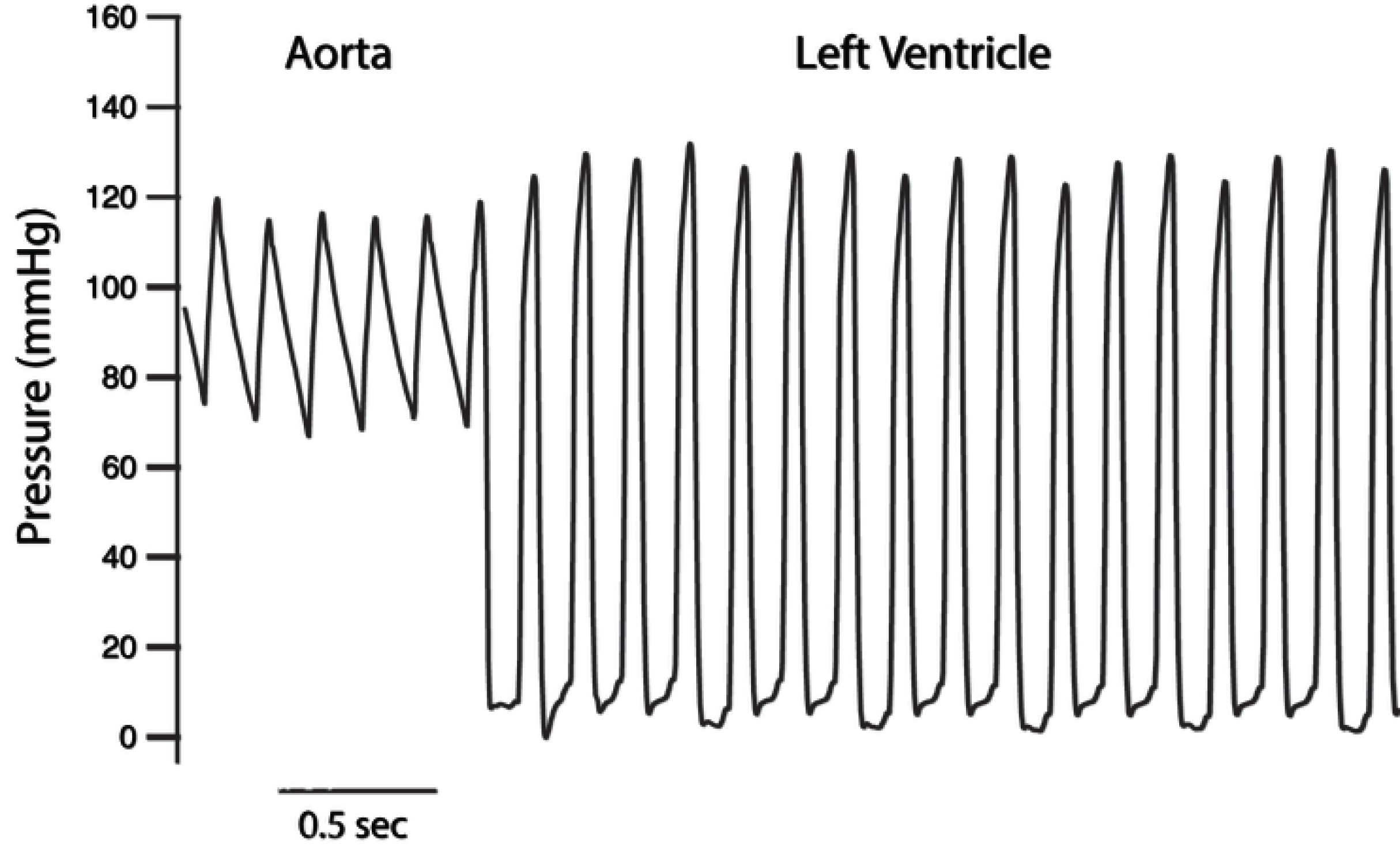
Invasive hemodynamics in control, HFpEF and AS)-E24 treated mice. Representative aortic and LV pressure tracing, and invasive hemodynamics were used to determine aortic pressure and LV function (see Table I).

**Table 1:** Invasive and Noninvasive Hemodynamic Parameters.

|  | **Control**<br>(n=4) | **HFpEF**<br>(n=4) | **ASO Rx**<br>(n=4) |
| --- | --- | --- | --- |
| HR (bpm) | 320±20 | 370±20 | 350±10 |
| Ao (mmHg) | 94±3/59±2* | 140±5/93±4 | 130±14/75±7^ |
| dP/dtmax (mmHg/s) | 6900±500 | 7300±1000 | 8500±800 |
| dP/dtmin (mmHg/s) | -8000±300* | -6000±500 | -8500±700* |
| LVEDP (mmHg) | 10±1* | 20±4 | 14±3 |
| τ (ms) | 8±1 | 10±1 | 8±0.4 |

***Noninvasive Hemodynamic Parameters***
|  | **Control**<br>(n=9) | **HFpEF**<br>(n=16) | **ASO Rx**<br>(n=14) |
| --- | --- | --- | --- |
| EF (%) | 60±2 | 60±1 | 60±2 |
| E (m/s) | 0.60±0.05* | 0.42±0.06 | 0.64±0.08* |
| A (m/s) | 0.38±0.03* | 0.24±0.02 | 0.47±0.04* |
| LV Septum (mm) | 0.72±0.02 | 0.73±0.01 | 0.78±0.02*^ |
| LV Posterior Wall (mm) | 0.73±0.04 | 0.75±0.02 | 0.81±0.03* |
| Lung (g) | 167±5* | 176±7 | 161±4* |

Invasive and noninvasive hemodynamic parameters (mean±SEM) in control, HFpEF and ASO-E24 treated male mice. Significant differences ( $p<0.05$ ) are indicated; \*, $p<0.05$ vs HFpEF; ^, $p<0.05$ vs control.

### Noninvasive Hemodynamics

Echocardiograms of normal control (n=9), HFpEF mice (n=17) and ASO-E24 treated mice are displayed in figure 3 (Table 1). Compared to normal controls, HFpEF mice have a normal ejection fraction (60%±2% control v 60%±1% HFpEF, p>0.05), and both early diastolic LV filling velocity (E; 0.60m/s±0.05m/s v 0.42m/s±0.06m/s, p=0.013) and late LV diastolic filing velocity (A; 0.38m/s±0.03m/s v 0.24m/s±0.05m/s, p=0.031) are depressed. Both septal (0.72mm±0.04mm v 0.73mm±0.01mm, p>0.05) and posterior wall (0.73mm±04mm v 0.75mm±0.02mm, p>0.05) thickness are similar in control vs HFpEF mice. Additionally, although body weight is similar in normal and HFpEF mice (31g±2g v 33g±1g, p>0.05), lung weight is significantly elevated (p=0.032) in the HFpEF mice (176mg±7mg, n=18) compared to normal control mice (167mg±5mg, n=6).

**Figure 3:**
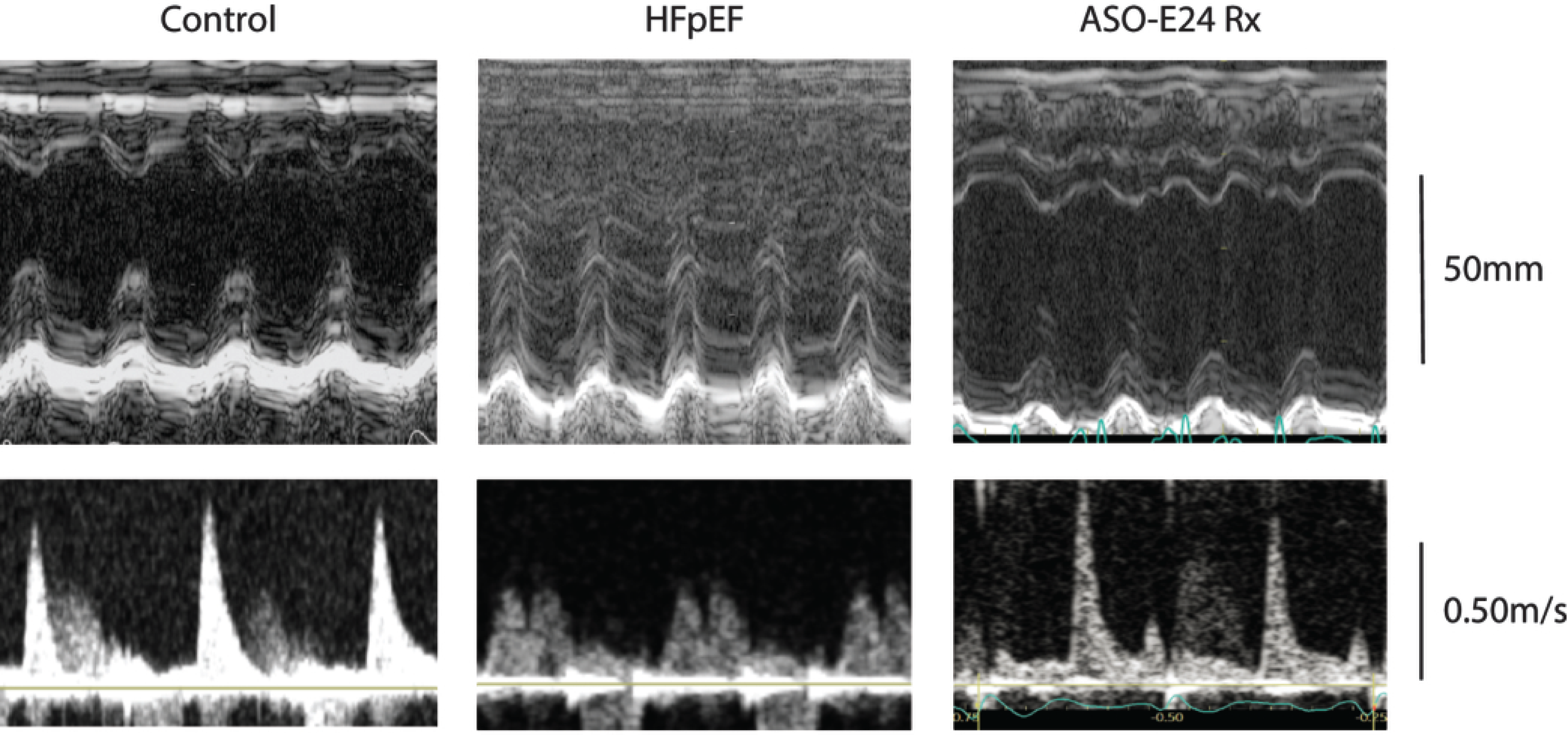
Echocardiograms of control, HFpEF and ASO-E24 treated mice. M-mode echocardiography and mitral doppler inflow velocity was used to assess LV hemodynamic parameters in control, HFpEF and ASO-E24 Rx mice (see Table I). Upper panels show M-mode images and lower panels are doppler signals of mitral inflow velocity.

ASO-E24 treated mice have a normal ejection fraction (59%±2%) and echocardiographic parameters of diastolic filling velocity are significantly higher than the mice with HFpEF; both E wave (0.64m/s±0.08m/s, p=0.034) and A wave (A, 0.47m/s±0.07m/s, p=0.016) are significantly different. Compared to mice with HFpEF, both body weight (29g±1g, p=0.009) and lung weight (161mg±4mg, p=0.030) are lower in the ASO-E24 treated mice. Additionally, septal (0.78mm±0.02mm, p=0.033) and posterior wall (0.81mm±0.03mm, p=0.38) thickness is greater in the ASO-E24 treated mice, but heart weight normalized to body weight is not different compared to HFpEF mice (4.3±0.2 v 4.6±0.2, p>0.05).

### MYPT1 E24

E24 exon inclusion expressed as E24 inclusion/(E24 inclusion+E24 exclusion) in control (n=7), HFpEF (n=7) and ASO-E24 Rx (n=7) mice is shown in Figure 4. E24 exon inclusion was higher in the mice with HFpEF (60%±6%) compared to normal controls (47%±7). ASO-E24 treatment significantly decreased E24 exon inclusion (22%±4%) compared to mice with HFpEF (p=0.001).

**Figure 4:**
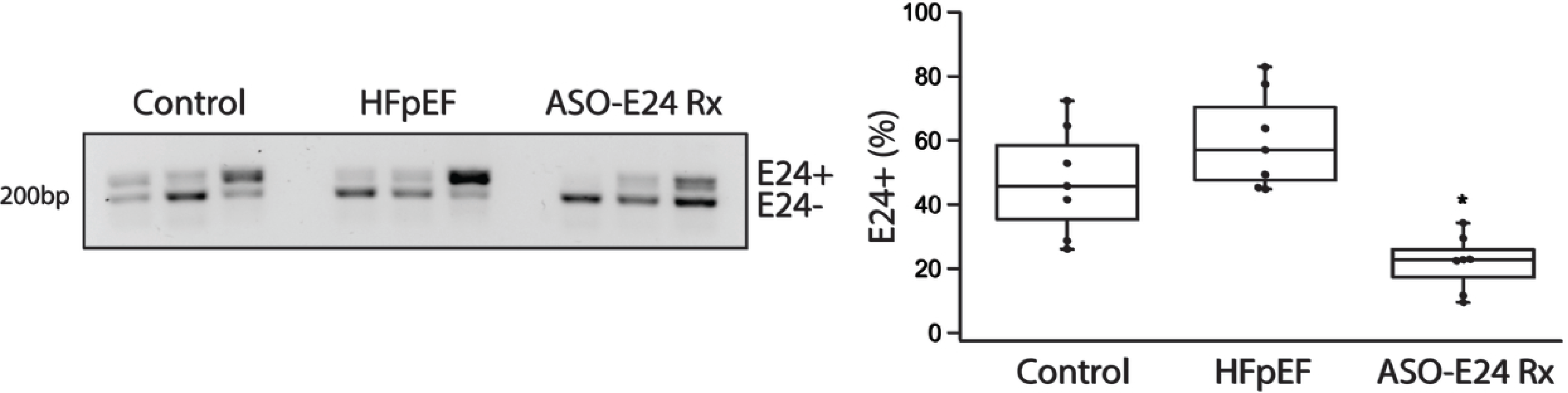
ASO-E24 treatment suppresses E24 exon inclusion in the MYPT1 transcript. RT-PCR demonstrates that E24 exon inclusion in control. HFpEF and ASO-E24 Rx mice. Data are summarized in the box plots (n=7 per group) with E24% representing (E24 inclusion/(E24inclusion+E24 exclusion))x100%; *, p<0.05 v HFpEF.

### MYPT1 LZ+ Expression

Alternative mRNA splicing of exon24 (E24) of the MYPT1 transcript produces MYPT1 isoforms that differ by the presence or absence of a COOH-terminal leucine zipper (LZ+/LZ-); E24 exclusion (E24-) produces an NO responsive LZ+ MYPT1, whereas E24 inclusion (E24+) produces an NO unresponsive MYPT1 ^20,21^. Thus, the increase in E24 inclusion in mice with HFpEF, as expected, results in a significant (p=0.001) decrease in the expression of the LZ+ MYPT1 isoform (1.0au±0.4au, n=6; Fig 5) compared to control mice (4.7au±0.7, n=6), and compared to HFpEF, ASO-E24 treatment increased LZ+ MYPT1 expression (2.0au±0.04au, n=6, p=0.001). Total MYPT1 expression (Fig 5) is also significantly (p=0.0001) lower in the HFpEF mice (1.0au±0.1au) compared to normal controls (4.7au±0.7au). Further, like LZ+ MYPT1 expression, ASO-E24 treatment increased total MYPT1 expression (2.0au±0.5au) compared to HFpEF (p=0.04).

**Figure 5:**
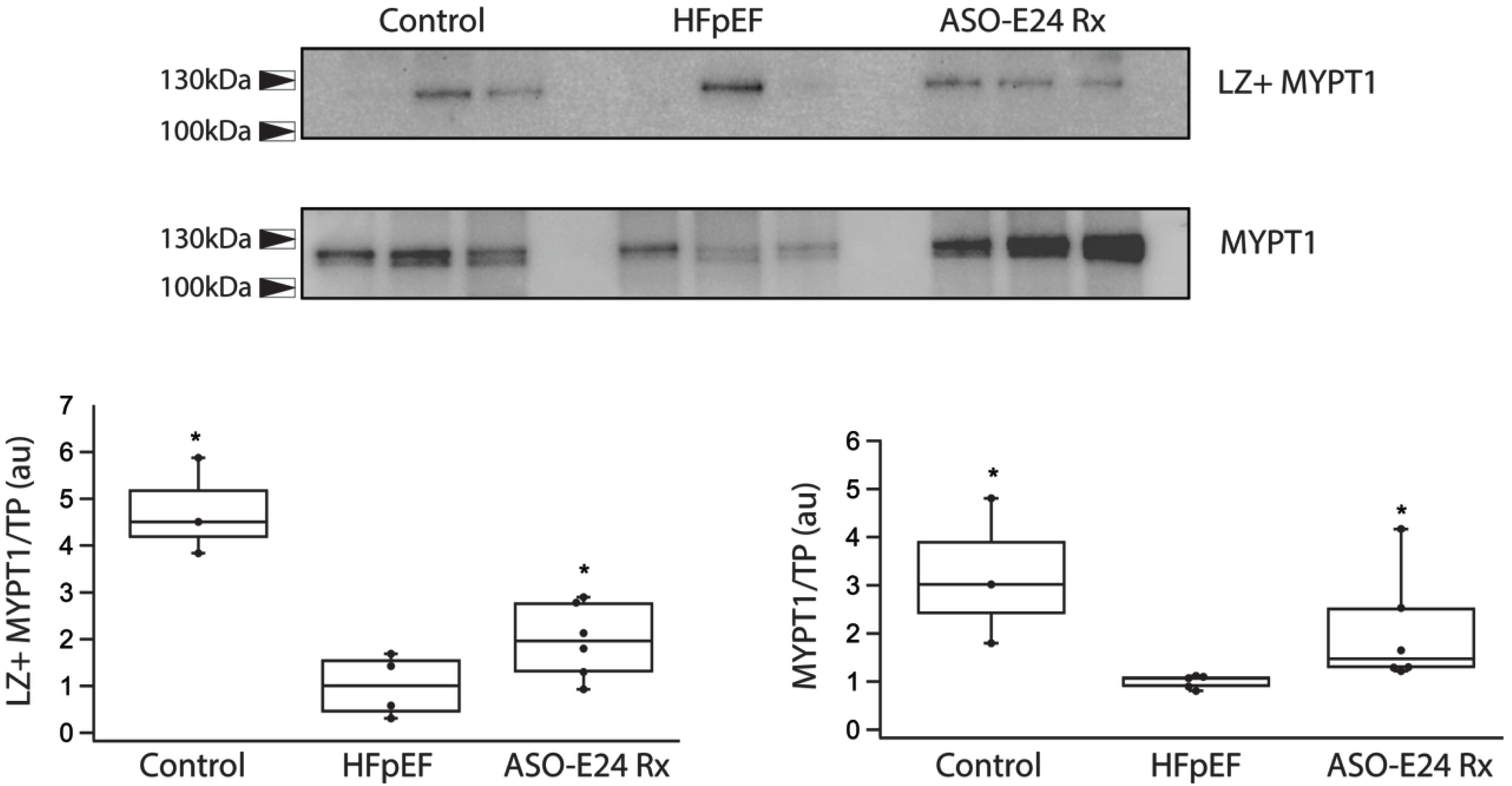
LZ+ MYPT1 expression is preserved with ASO-E24 treatment. Immunoblots demonstrate both total MYPT1 and LZ+ MYPT1 expression (n=6 per group) are lower in HFpEF than either control or ASO-E24 Rx. Data of MYPT1/TP and LZ+ MYPT1/TP are summarized in the box plots; *, p<0.05 v HFpEF.

### Cardiac Contractile Proteins

To determine whether changes in vascular reactivity alter cardiac protein expression and phosphorylation, we examined two key regulators of the intracellular Ca^2+^ transient during cardiac muscle relaxation, SERCA2 and PLN. SERCA2 expression was similar (p>0.05) in normal control (100au±2au, n=6) and HFpEF (110au±9au, n=6) animals, but ASO-E24 treatment reduced (44au±5au, n=6, p=0.007) the expression of SERCA2 (Fig 6). PLN expression was lower in control (100au±10au) compared to HFpEF (240au±50au, p=0.023) and ASO-E24 treatment lowered PLN expression to the level (140au±20au) control animals. The phosphorylation of PLN (p-PLN/PLN) is higher (p=0.010) in the mice with HFpEF (200au±20au) than control mice (110au±20au) and ASO-E24 Rx lowered PLN phosphorylation (180au±40au, p>0.05) to that of normal control mice (Fig 6).

**Figure 6:**
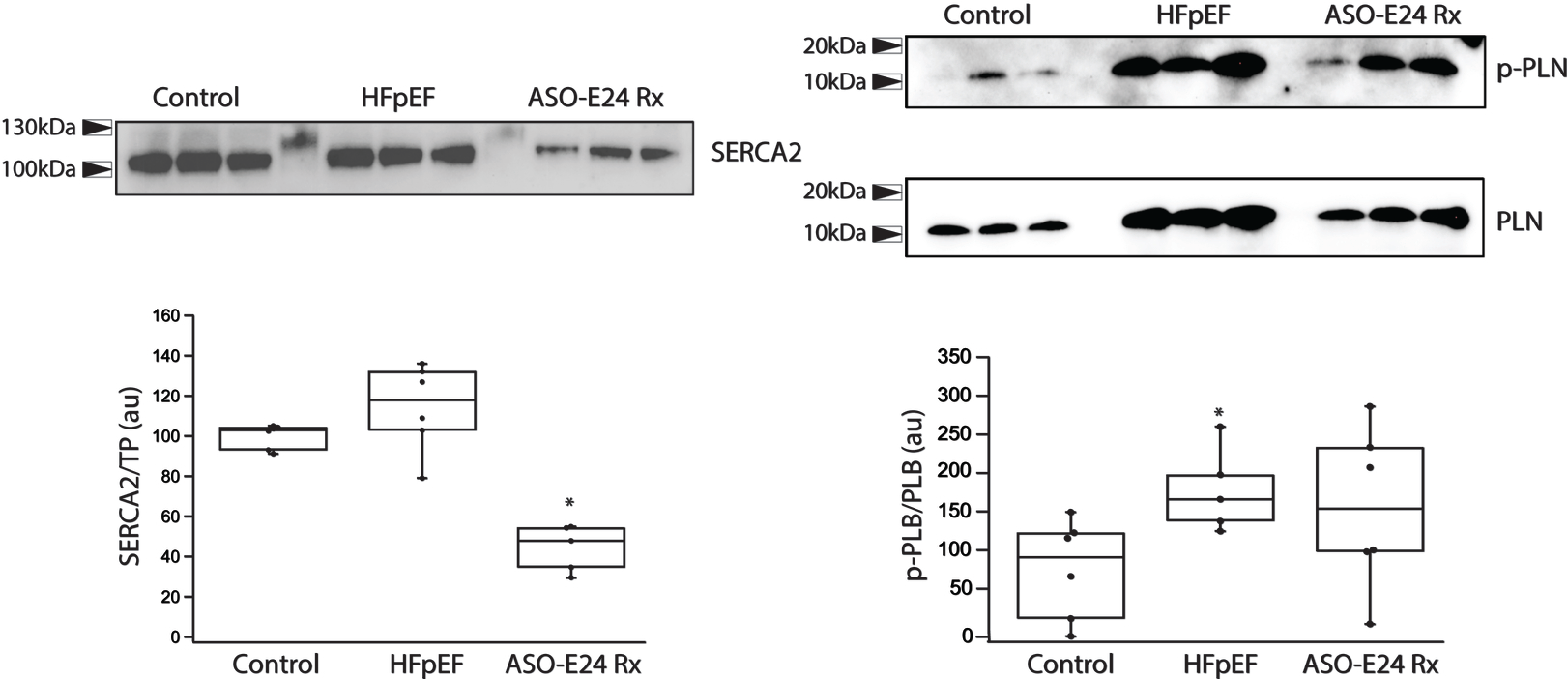
ASO-E24 treatment prevents an increase in PLN phosphorylation. Immunoblotting demonstrates that SERCA2 expression is lower in ASO-E24 Rx mice and PLN phosphorylation is higher in HFpEF mice (n=6 per group). Data of SERCA2/TP and p-PLN/PLN are summarized in the box plots; *; p<0.05 v Control.

Our results also demonstrate that the phosphorylation of TnI is similar (p>0.05) in control (44au±6au), HFpEF (60au±8au) and ASO-E24 (58au±7au) treated mice (Fig 7). However, the phosphorylation of MyBP-C (Fig 7) is higher in mice with HFpEF (197au±10au) than either control (150au±21au, p=0.034) or ASO-E24 treated mice (160au±14au, p=0.030).

**Figure 7:**
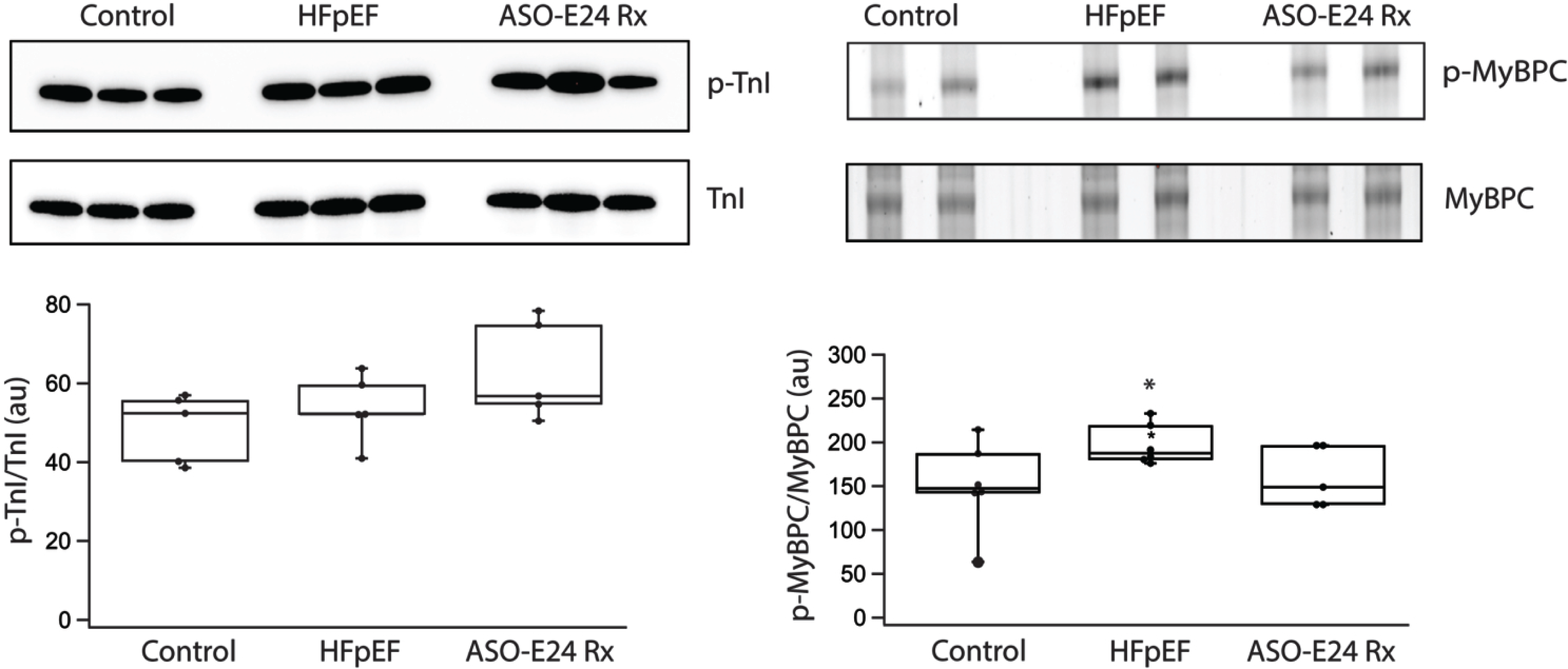
ASO-E24 treatment prevents the increase in MyBP-C phosphorylation. Immunoblotting demonstrates that the phosphorylation of TnI is similar in control, HFpEF and ASO-E24 Rx mice (n=6 per group). Pro-Q Diamond staining (upper) and SYPRO-Ruby staining (bottom) were used for visualization of p-MyBP-C and MyBP-C, respectively. Data of Data of p-TnI/TnI are summarized in the box plots p-MyBP-C/MyBP-C are summarized in the box plots; *, p<0.05 v HFpEF.

### Vascular Reactivity

Vascular reactivity was assessed by determining the magnitude of the relaxation to 8Br-cGMP. The contractile response of mesenteric arteries to KCl depolarization is phasic with a rapid rise in force to a peak and then fall to a sustained plateau (Fig 8). Peak force in control (8.3mN/mm±1.1mN/mm^2^, n=6), HFpEF (8.5mN/mm^2^±1.1mN/mm^2^, n=6) and ASO treated mice (7.4mN/mm^2^±1.3mN/mm^2^, n=6) as well as sustained force in control (4.9mN/mm^2^±1.2mN/mm^2^), HFpEF (1.0mN/mm±0.4mN/mm^2^) and ASO treated mice (0.8mN/mm^2^±0.21mN/mm^2^mN) are not different. However, the relaxation to 100µM 8BrcGMP is significantly larger in control (65%±5%, p=0.0307) and ASO treated mice (72%±9%, p=0.0270) than in the HFpEF mice (44%±9%).

**Figure 8:**
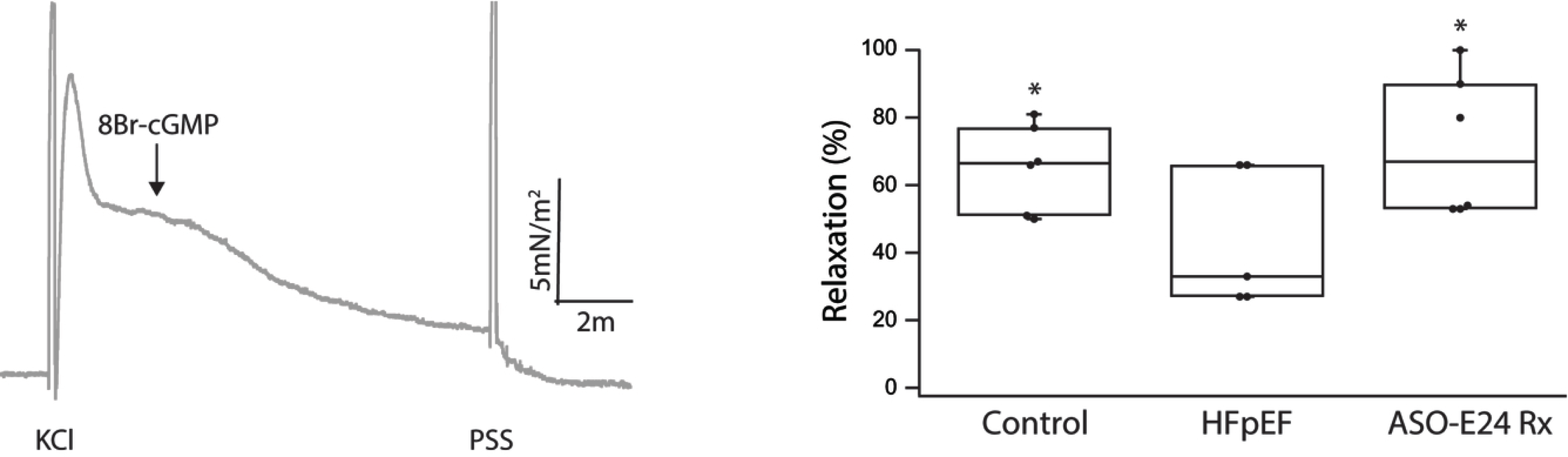
ASO-E24 treatment maintains normal vascular reactivity. Force in response to KCl is phasic and 8Br-cGMP produces relaxation. The magnitude of relaxation in control, HFpEF and ASO-E24 treated mice (n=6 per group). Relaxation data are summarized in the box plots; *, p<0.05 v HFpEF.

### Cardiac Hypertrophy

Myocyte CSA is significantly higher in the HFpEF mice (619.3±49.8 µm^2^, n=6, Fig 9) than in control (375.7±46.7µm^2^, n=6, p=0.0026) and ASO treated mice (440.5±39.4µm^2^, n=6, p=0.0010).

**Figure 9.**
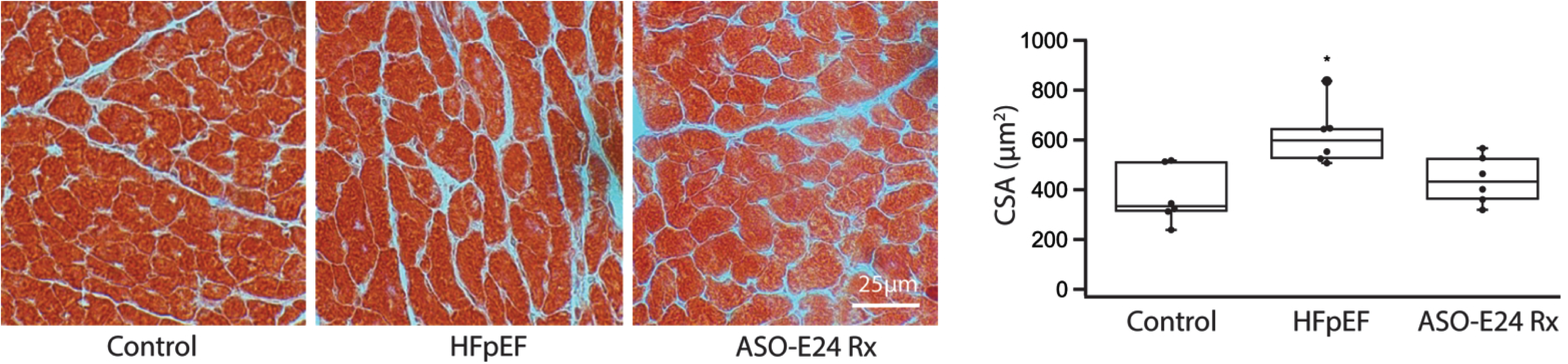
ASO-E24 treatment preserves myocyte cross-sectional area (CSA). Representative hematoxylin–eosin (H&E)–stained cardiac tissue sections are shown; scale bar, 25 μm (applies to all images). CSA values are presented as box plots, n=6 per group; *p < 0.05 vs. Control and ASO-E24 Rx.

## Discussion

Male mice fed a HFD and L-NAME in the drinking water for 4 weeks had a normal EF, a higher LVEDP, diastolic dysfunction and increased lung weight compared to normal control mice (Table I). These findings are consistent with previous reports ^33,35^, and demonstrate that male mice treated with a HFD and L-NAME develop HFpEF.

The pathological cascade that leads to HFpEF is thought to begin with abnormal vascular reactivity, including a decrease in the sensitivity of the vasculature to NO mediated vasodilatation and coupled with subsequent changes in cardiac contractility and energetics ultimately producing HFpEF ^13,14^. Vascular tone is regulated by the phosphorylation of the regulatory light chain (RLC) of smooth muscle (SM) myosin, which is controlled by the activity of myosin light chain (MLC) kinase and MLC phosphatase ^15^. MLC phosphatase is a trimeric enzyme consisting of a catalytic, myosin targeting (MYPT1) and 20kDa subunits ^15,18,19^. Alternative mRNA splicing of E24 of the MYPT1 transcript produces two distinct MYPT1 isoforms; E24 exclusion produces an NO responsive LZ+ MYPT1, while E24 inclusion produces an NO unresponsive LZ- MYPT1 ^20,21^. Further, LZ+/LZ- MYPT1 isoform expression has been demonstrated to regulate the sensitivity of smooth muscle to NO mediated vasodilation ^22–25^. This study used ASO-E24 (Fig 1) to suppress E24 exon inclusion in the MYPT1 transcript ^32^ and increase the expression of the LZ+ MYPT1 isoform ^32^, which would enhance vascular sensitivity to NO mediated vasodilation ^22–25^. Since LZ+ MYPT1 expression has been shown to decrease in HFpEF ^30,31^, preventing the decrease in LZ+ MYPT1 expression could interrupt the pathological cascade and prevent the development of HFpEF. In ASO-E24 treated mice, hemodynamics showed that -dP/dtmin and LVEDP were not different compared to normal controls while echocardiography demonstrated that the ASO-E24 mice are hypertensive and develop LV hypertrophy, but other hemodynamic parameters including EF, LV diastolic properties and LVEDP are normal as is lung weight (Table I). Further, myocyte cross-sectional area was preserved in the ASO-E24 treated mice. Thus, these data show that ASO-E24 treatment prevents the development of HFpEF in response to a HFD and L-NAME.

We have previously demonstrated that LZ+ MYPT1 expression is lower in both rats ^44^ and humans with HFpEF ^31^. Similar to these previous studies, our data demonstrate a reduction in LZ+ MYPT1 (Fig 5) in HFpEF mice. Our previous publications demonstrate a decrease in LZ+ MYPT1 expression produces a decrease in the sensitivity to NO mediated vasodilatation ^15,22–25^, which has been suggested to represent the initial step in the pathological cascade leading to HFpEF ^13,14^. Additionally, our data demonstrate that total MYPT1 expression is also lower in animals with HFpEF (Fig 5), and a decrease in total MYPT1 expression would increase vascular tone ^15^, which also contributes to abnormal vascular reactivity, which is hypothesized to be a factor in the initial step in the pathology that produces HFpEF ^13,14^. Consistent with previous reports ^32^, ASO-E24 treatment suppresses E24 exon inclusion in the MYPT1 transcript (Fig 4) and increases the expression of the LZ+ MYPT1 isoform in smooth muscle (Fig 5). Thus, our results show that MYPT1 and LZ+ MYPT1 expression are reduced in animals with HFpEF and ASO-E24 treatment suppresses E24 exon inclusion in the MYPT1 transcript and prevents the decrease in LZ+ MYPT1 and total MYPT1 expression observed in mice with HFpEF (Figs 4&5). Our data demonstrate that HFpEF mice also have reduced vascular reactivity represented by a decrease in 8Br-cGMP mediated force relaxation. Additionally, ASO-E24 treatment, which maintains LZ+ MYPT1 expression (Fig 5) also maintains a normal relaxation to 8Br-cGMP. We have demonstrated that LZ+ MYPT1 expression controls the sensitivity to NO mediated vasodilatation ^15,22–25^, and thus consistent with these previous results, ASO-E24 treatment maintains normal LZ+ MYPT1 expression and vascular reactivity. Further, ASO-E24 Rx mice do not have HFpEF (Table I), which is consistent with the hypothesis that the pathological cascade that leads to the development of HFpEF begins with abnormal vascular reactivity, specifically a decrease in LZ+ MYPT1 and total MYPT1 expression. Further, our data demonstrate that maintaining normal vascular reactivity prevents the development of HFpEF. Others have demonstrated that mice treated with a HFD and L-NAME (HFpEF) are hypertensive ^33^, which is consistent with our data (Table 1). Increasing LZ+ MYPT1 expression in vascular smooth muscle has been shown to increase the sensitivity to NO mediated vasodilatation ^22–25^ and blood pressure is lower in transgenic mice with increased LZ+ MYPT1 expression ^20,26^. However, the blood pressure in the ASO-E24 treated and HFpEF mice is not different (p>0.05, Table1), which demonstrates that a decrease in blood pressure does not mediate the beneficial effect of ASO-E24 treatment. Further, antihypertensive therapy has not been demonstrated to be effective in treating HFpEF ^5,6,47,48^. These data suggest the prevention of HFpEF due to ASO-E24 treatment is mediated by maintaining LZ+ MYPT1 expression and preserving normal vascular reactivity.

In a rodent HFrEF model, both ACE inhibition ^27^ and ARB treatment ^49^, but not prazosin ^27^, have been shown to maintain normal LZ+ MYPT1 expression and improve EF. Similar to our results in HFpEF, these data suggest that preserving normal vascular reactivity could underlie some of the therapeutic benefits of both ACE inhibition and ARB treatment in HFrEF ^3,50^. Moreover, LZ+ MYPT1 expression is reduced in mice with HFpEF (this study), rats with HFpEF ^30^ and humans with HFpEF ^31^. The fall of LZ+ MYPT1 expression would decrease sensitivity of the vasculature to NO/cGMP/PKG signaling (Fig 8, ^15,22–25^) and could explain the lack of clinical benefit of HFpEF trials aimed at enhancing PKG signaling ^5–7^.

Similar to HFrEF in rodents produced by LAD ligation ^27,49^, the present study demonstrates that maintaining normal vascular reactivity prevents the development of HFpEF. However, HFpEF is also associated with diastolic dysfunction ^10–12^ and the rate of decline of the intracellular Ca^2+^ transient is an important determinant of cardiac relaxation ^51^. SERCA2 and PLN regulate the rate of decline of the Ca^2+^ transient in cardiomyocytes ^51^. PLN decreases the Ca^2+^ affinity of SERCA2 and phosphorylation of PLN relieves the inhibition of SERCA2 ^51^. In rats with HFpEF, both SERCA2 expression and PLN phosphorylation are lower ^30^, suggesting that a decrease in the rate of decline of the Ca^2+^ transient contributes to diastolic dysfunction in the rat HFpEF model. Our data show that SERCA2 expression is similar in control and HFpEF mice, but lower in the ASO-E24Rx mice (Fig 6). Further, PLN expression and phosphorylation are higher in mice with HFpEF (Fig 6). The data in the present study suggest that in mice, diastolic dysfunction in HFpEF is not due to a change in the Ca^2+^ transient, but rather possibly due to increased ventricular stiffness ^52^, driven in part by higher collagen content ^30^. If this is the case, the increase in PLN expression and phosphorylation observed in the mice with HFpEF could represent a compensatory mechanism to normalize and/or increase the rate the decline of the Ca^2+^ transient and cardiomyocyte relaxation. Mice treated with ASO-E24 have normal diastolic properties (Table I). The ASO-E24 mediated normalization of diastolic function may have alleviated the compensatory increase in SERCA2 expression and normalization of PLN expression and phosphorylation. Although TnI phosphorylation was not altered in HFpEF (Fig 7), MyBP-C phosphorylation increased in HFpEF and ASO-E24 treatment decreased the phosphorylation of MyBP-C (Fig 7). MyBP-C stabilizes the myosin SRX state and phosphorylation of MyBP-C moves myosin S1 heads away from the thick filament backbone to interact with actin ^53–55^ and increases the AMATPase rate ^53,56^. Further, phosphorylation of MyBP-C has been demonstrated to increase actin sliding velocity ^57^, force redevelopment and enhance diastolic function ^58^. These data would suggest that the increase in MyBP-C phosphorylation observed during HFpEF is a compensatory response to improve diastolic dysfunction. Thus, the improvement of diastolic properties with ASO-E24 treatment could prevent the increase in MyBP-C phosphorylation observed in the animals with HFpEF.

In contrast to our data demonstrating similar TnI phosphorylation in HFpEF and increased MyBP-C phosphorylation in HFpEF, are the recent data reported in patients with HFpEF ^59^. This study used samples of RV cardiac muscle and divided HFpEF patients into two groups based on BMI; obese (BMI, 31kg/m^2^±6kg/m^2^) and very obese (BMI, 43kg/m^2^±8kg/m^2^). Compared to normal controls, phosphorylation of MyBPC at PKA sites, TnI (S23, S24) and (S275, S284, S304), was reduced in both groups. However, total phosphorylation of MyBP-C was reduced only in the obese, but not very obese patients with HFpEF, while total TnI phosphorylation was reduced in both HFpEF groups. However, we used cardiac muscle from the LV and the human study from the RV ^59^, and the weights of our control (31g±2g) and HFpEF (33g±2g) mice were similar (p>0.05). Protein phosphorylation may be different in LV vs RV and the lack of a significant weight difference also could contribute to differences in TnI and MyBP-C phosphorylation in LV cardiac samples in our HFpEF mice compared to samples from RV biopsies in obese and very obese patients ^59^.

## Conclusion

These findings suggest that ASO-E24 treatment not only prevents the decrease in LZ+ MYPT1 expression, but maintaining normal LZ+ expression and vascular reactivity prevents diastolic dysfunction and/or changes in cardiac contractility that contribute to the pathogenesis of HFpEF. These data are consistent with the hypothesis that a decrease in NO mediated vasodilatation, or abnormal vascular reactivity, is the initial step in the pathological cascade that leads to HFpEF ^13,14^. Furthermore, our data indicate that preserving LZ+ MYPT1 expression and normal vascular reactivity prevents the subsequent changes in protein expression and phosphorylation that regulate cardiac contractility (Fig 6&7) and myocardial energetics that produce HFpEF. Thus, rather than treating abnormal vascular reactivity with vasodilators, maintaining normal vascular reactivity may represent a novel therapeutic strategy for HFpEF.

Our data demonstrate that suppressing E24 exon inclusion in the MYPT1 transcript with ASO-E24 maintains LZ+ MYPT1 expression and prevents the development of HFpEF. Further, the 50bp sequences flanking E24 are highly conserved across mammals ^60^, and thus, our data suggest that ASO-mediated suppression of E24 exon inclusion may represent a novel therapy for HFpEF.

## Acknowledgements

This study was supported by a Mayo Clinic CV Prospective Research Award (FVB).

## Data Availability

Data are available on request.

## Declaration of generative AI and AI-assisted technologies in the manuscript preparation process

AI was not used in the preparation of the manuscript.

## Author Contributions

YSH performed the experiments, analyzed the data and wrote the manuscript; TMP performed and analyzed the echocardiograms; BZ performed invasive hemodynamics; MJF measured myocyte CSA; GCS participated in revising the manuscript; FVB designed the study, supervised the experiments and data analysis and revised the manuscript.

